# Fraction and Spatial Organization of Tumor-Infiltrating Lymphocytes Correlating with Bladder Cancer Patient Survival

**DOI:** 10.1101/666503

**Authors:** Hongming Xu, Sunho Park, Tae Hyun Hwang

## Abstract

Recent studies based on deep learning on pathology images have shown the potential prognostic role of Tumor-infiltrating lymphocytes (TILs) in many cancer types. In general, most of these studies focused on detecting, quantifying and identifying TILs and spatial organization in entire whole slide images (WSIs) including both non-cancer and cancer regions. In this work, we hypothesize that TILs present in cancer regions not entire WSIs could have prognostic values. Here we present an automatic technique based on convolutional neural networks (CNN) that detects both cancer and TILs regions, and explores fraction and spatial organization of TILs in the detected cancer region from WSIs. A novel set of quantitative histological image features reflecting TILs fraction and spatial organization is extracted and used to stratify patients with distinct survival outcomes. In experiments using 277 different patient pathology slides selected from The Cancer Genome Atlas (TCGA) bladder cancer cohort, we show that bladder cancer patient survival is significantly correlated with our designed TIL-related image features, with a log-rank test *p* value below 0.05. Our method is extensible to histopathology images of other organs for predicting patient survival via TIL analysis.

## 1 Introduction

Bladder cancer ranks the fourth most common cancer in men and is a world-wide leading cancer death among both men and women [1]. To reduce patient mortality from bladder cancer, it is critical to determine appropriate strategies for disease treatment or monitoring based on individual’s predicted risk. Genetic and biological markers have been examined extensively for their potential to signal progression or risk of disease. However, one key challenge is that not all patients would have adequate tumor tissues excised for genetic or biological testing. In addition, genetic testing (e.g., DNA sequencing) is usually expensive and time consuming, which is difficult to be incorporated into public health practice. On the other hand, hematoxylin-eosin (HE) stained tissue diagnostic slides are virtually available for all bladder cancer patients. It would be promising to explore effective markers in histological images to predict bladder cancer patients into different risk groups.

Tumor-infiltrating lymphocytes (TILs) are lymphocytes of the host immune system that have been observed within tumor sites [1]. Presumably lymphocytes migrate to tumor regions to combat the growing malignant cells. The density and distribution of TILs have been shown with prognostic significance in many cancers [1]. Recently, there have been a few studies that attempt to characterize TIL-related histological image features and correlate them with clinical outcomes such as patient survival or tumor recurrence. Corredor et al. [3] presented a study that predicts likelihood of recurrence in early-stage non-small cell lung cancer by using TIL-related features computed in tissue microarrays. They first construct spatial graphs based on lymphocytes and non-lymphocytes distinction. Two sets of image features describing spatial distribution of lymphocytes and relationship between lymphocytes and non-lymphocytes are extracted. The quadratic discriminant analysis is used to predict patients into one of two classes: recurrence or non-recurrence. Saltz et al. [2] presented a study to generate TIL maps across 13 TCGA tumor types. Due to the huge size of the WSI, it is a computational challenge to detect lymphocytes separately as done in [3]. To overcome computational limitations, this study divides the WSI into many non-overlapping blocks and trains CNN models to predict image blocks as lymphocyte patches or not. Affinity propagation is applied to cluster predicted lymphocyte patches. The clustering index (e.g., C index) is used to correlate with patient survival and tumor subtypes. Although the clustering index is shown to be informative, this study does not distinguish tumor and non-tumor regions in WSIs and thus cannot effectively describe densities and distributions of lymphocytes within tumor regions. In addition, it is worth to note that this study did not find associations of TIL local spatial structure and survival in bladder cancer.

This study is extended from the reference [2], where TIL-related histological image features are extracted from bladder cancer WSIs. The major contributions for this study include: (1) we design a novel set of histological image features that reflect fraction and spatial organization of TIL regions in WSIs; (2) The designed TIL-related image features are found to be significantly correlated with TCGA bladder cancer patient survival by using the consensus clustering method.

## 2 Methods

Fig. 1 shows the pipeline of presented technique. As observed in Fig. 1, the technique has four modules: tumor detection, TILs detection, TILs characterization and consensus clustering. The details of four modules are described below.

**Fig. 1:**
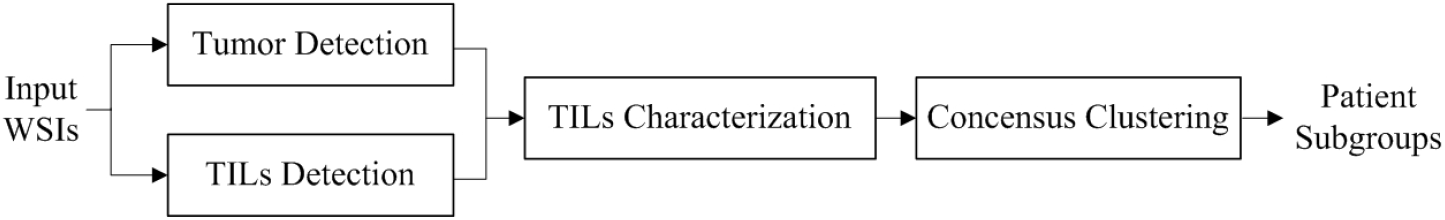
Pipeline of the presented technique.

### 2.1 Tumor Detection

The WSI usually has both tumor and non-tumor regions. In order to detect tumor regions, we trained a deep learning model which has a similar architecture with the VGG16 model [4]. Due to the relatively less number of training samples, our trained model has much less number of trainable parameters (about 0.28 million) compared to the VGG16 model. Fig. 2 shows our deep learning architecture for tumor detection. As shown in Fig. 2, the model has 13 convolutional layers with either 8, 16, 32 or 64 feature map channels. The model has 4 3×3 max-pooling layers, 2 dense layers (with 128 neurons), 2 drop-out layers (with 0.25 drop-out rate). The output layer uses the Sigmoid activation function.

**Fig. 2:**
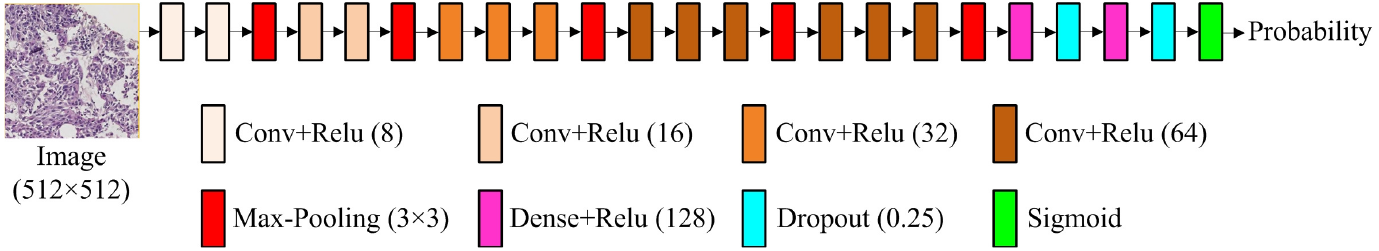
Deep learning architecture for tumor detection.

To train the model, a pathologist was asked to annotate tumor regions in 60 WSIs. The annotated WSIs were then divided into 512×512 image blocks (at 20× magnification), which included about 20,000 tumor and 22,000 non-tumor blocks. Image augmentations including rotation, zooming, flipping and color-based augmentations were applied along with training. The RMSprop optimizer with default parameter settings in Keras deep learning package was used to minimize binary cross entropy loss function. The maximum training epoch was set as 30. The batch size was set as 128. The early stopping was applied if there was no performance improvement on validation set. The trained model is used to detect cancer regions in the testing WSI, which outputs a probability map. An empirically selected threshold of 0.5 is applied on the probability map to binarize tumor regions. Small holes of the tumor mask *I*_*t*_ are filled by morphological operations. Fig. 3(a) shows a bladder caner WSI overlapped with heat map of tumor detection. Fig. 3(b) shows the probability map of tumor regions. Fig. 3(c) illustrates the final tumor detection result highlighted with red contours.

**Fig. 3:**
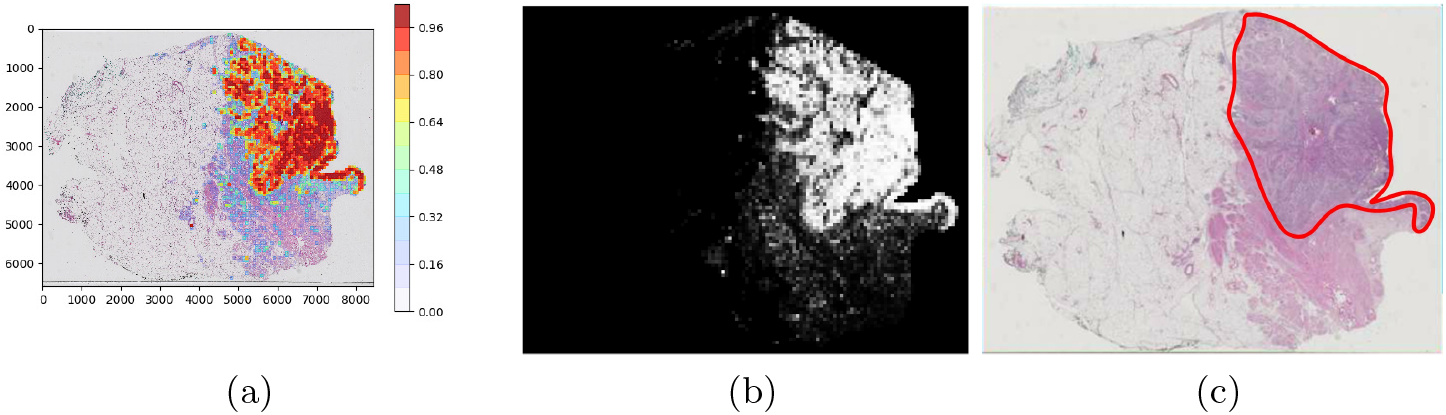
Example of tumor detection. (a) WSI overlapped with head map of tumor detection. (b) Probability map of tumor regions. (c) Tumor detection result.

### 2.2 TILs Detection

In parallel with tumor detection, TILs detection is performed by using two pre-trained convolutional neural network (CNN) models: lymphocyte CNN and necrosis segmentation CNN [2]. The lymphocyte CNN is a 18-layer network with 14 convolutional layers, 3 pooling layers and 1 dense layer. The lymphocyte CNN classifies image blocks (100×100 pixels) of WSI at 20× magnification into those with lymphocyte infiltration and those without. Since necrotic regions are usually recognized as lymphocyte-infiltrated regions because of similar nuclei characteristics, the necrosis segmentation CNN is then applied to eliminate false positives from the lymphocyte CNN. The necrosis segmentation CNN has the same architecture as the DeconvNet [6], which is performed on image patches of 333×333 pixels (about 7× magnification) and outputs pixel-wise segmentation result. The recognized lymphocyte-infiltrated patches belonging to necrotic regions are finally classified as non-lymphocyte-infiltrated. For more details about lymphocyte CNN and necrosis segmentation CNN, please refer to [2, 6]. Fig. 4 shows TILs detection, where red pixels indicate lymphocyte-infiltrated regions.

**Fig. 4:**
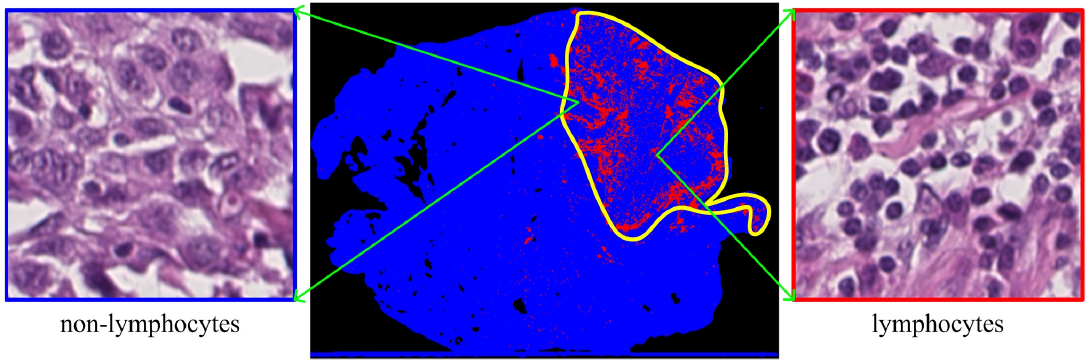
Illustration of TILs detection, where red pixels highlight regions with lymphocytes and yellow contours highlight tumor regions.

### 2.3 TILs Characterization

Based on tumor and TILs detection, we design quantitative features to measure fraction and spatial organization of lymphocytes within tumor regions.

#### TILs fraction

To capture fraction of TILs, we compute 3 histological image features as follows:

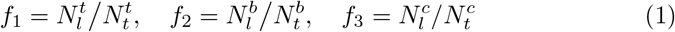

where 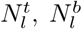 and 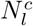 represent the number of lymphocyte pixels within the tumor region (see red pixels in Fig. 5(a)), tumor border region and tumor center region, respectively. 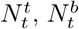 and 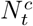 represent the number of pixels belonging to tumor region, tumor border and tumor center regions, respectively. To obtain the tumor border region, we first erode the tumor mask *I*_*t*_ (obtained by tumor detection) by a disk structuring element with a radius of 5 to obtain a shrunk tumor mask *I*_*s*_. We then dilate the mask *I*_*t*_ by the same disk structuring element to obtain a dilated mask *I*_*d*_. The tumor border mask is determined by applying exclusive OR operation on masks *I*_*s*_ and *I*_*d*_. Fig. 5(b) illustrates the corresponding tumor border region. To obtain the tumor center region, the iterative erosion operation [5] is applied on tumor mask *I*_*t*_ to generate a tumor center mask *I*_*c*_, where the number of tumor pixels within *I*_*c*_ is half of that within *I*_*t*_. Fig. 5(c) illustrates the corresponding tumor center region.

**Fig. 5:**
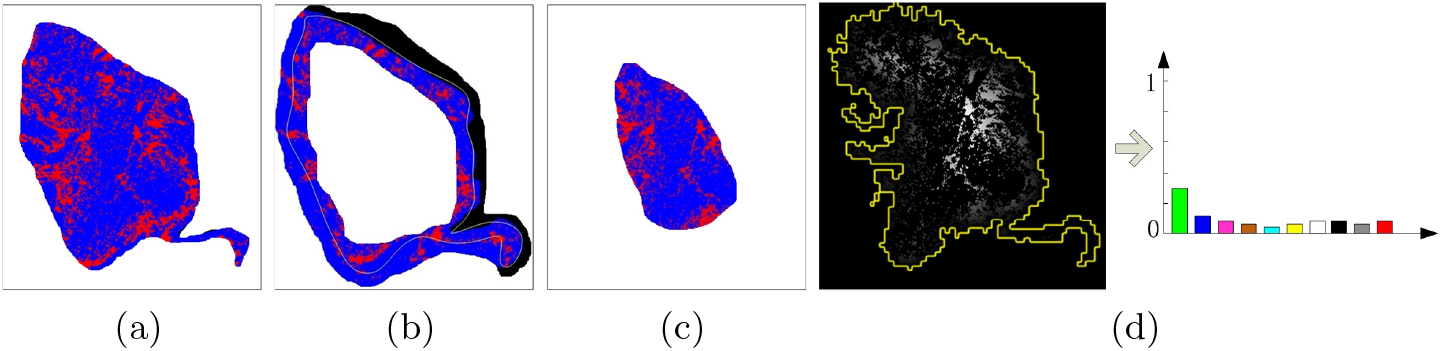
Illustration of TILs characterization. Fraction of TILs within (a) tumor region, (b) tumor border region, (c) tumor center region. (d) Normalized distance map of TIL pixels and histogram features.

#### TILs spatial organization

To capture the spatial organization of TILs, we first compute the normalized nearest distance to the non-tumor region for every TIL pixel, i.e., 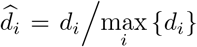, 1 ≤ *i* ≤ *n*, where *d*_*i*_ is the nearest distance from the *ith* lymphocyte pixel to the non-tumor region and *n* is the number of TIL pixels. We then divide elements 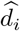 into a number of equally spaced bins (i.e., 10 bins in this study). Finally, we count the number of elements within each bin and normalize it as the quantitative feature, which is computed as:

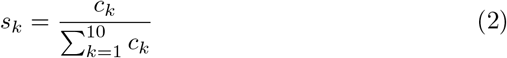

where *c*_*k*_ represent the number of TIL pixels within the *kth* bin, 1 ≤ *k* ≤ 10. Fig. 5(d) illustrates histogram features extracted from the normalized distance map of TIL pixels. The features *s*_*k*_ capture spatial organization of TILs by statistically computing distances from TIL pixels to the nearest non-tumor regions.

After TILs characterization, each patient WSI corresponds to a 13 dimensional TIL-related feature vector.

### 2.4 Consensus Clustering

To stratify patients into different prognosis groups e.g., good vs poor (vs intermediate), we apply consensus clustering on the TIL-related feature vectors. Consensus clustering, also known as cluster ensemble, gives a single consolidated clustering result by aggregating results from multiple runs of clustering algorithms. In this study, we adopt a variant of non-negative matrix factorization (NMF) [8] as the clustering algorithm and follow aggregation method in [7] to combine multiple clustering results. By setting the number of subgroups *k* to be 2 or 3, we conduct consensus clustering by three steps. (1) we run the NMF method (with the latent dimension set as *k*) 1000 times on randomly selected 80% of the samples. (2) we combine these 1000 different clustering results by constructing a consensus matrix **M** (samples×samples), where each element of **M** has a value equal to the time of two corresponding samples appeared in the same cluster dividing the total time of two samples included for experiments. Each element in **M** can be considered as the probability of two corresponding samples assigned to the same cluster. (3) the final clustering result is obtained by applying hierarchical clustering on matrix **M**. The input to hierarchical clustering are distances between samples, i.e., the distance between the *i* and *j* samples is computed as 1 − *M*_*ij*_, where *M*_*ij*_ is the (*i, j*) element of matrix **M**.

## 3 Experimental Results

### 3.1 Dataset Description

A cohort of 386 bladder cancer patients with diagnostic slides and survival information was downloaded from TCGA data portal (project TCGA-BLCA). Among 386 patients, 277 patients were selected for survival stratification, while the remaining patients were excluded due to either lack of survival information or with poor quality slides (e.g., pen markers). Although there are a small number of patients with more than 1 diagnostic slides, for simplicity we only select the first diagnostic slide with DX1 suffix from every patient. Each digitized slide has a maximum scanning resolution either at 20× or 40× (0.25*μm* per pixel).

### 3.2 Patient Survival Stratification

We conducted patient stratification by using different feature sets including local binary patterns (LBP) [9], transfer learning on Xception (TLX) [10], TIL fraction (TIL-F), TIL spatial organization (TIL-S) and TIL fraction with spatial organization (TIL-F-S). For LBP and TLX, the features describing the WSI were extracted by three steps. (1) the detected tumor regions (at 20×) were divided into many non-overlapping blocks (1024×1024). (2) LBP or transfer learning features were separately computed from all image blocks. (3) the mean of features from all blocks was computed to represent the WSI. We computed multi-scale LBP by encoding 16 neighboring pixels which resulted in a 72 dimensional feature vector. Transfer learning on Xception model resulted in a 2048 dimensional feature vector for the WSI. Due to the high feature dimensionality, we also tested the principal component analysis (PCA) to reduce feature dimensions from 2048 to 100 (explained 98% of variance) for TLX. Using these feature sets, we separately divided patients into *k* = 2 (or 3) prognosis groups, i.e., good vs poor (vs intermediate). For each *k* and feature set, we calculated a *p*-value from the log-rank test which tests the null hypothesis that there is no difference in overall survival between independent groups.

Table 1 shows the log-rank test *p*-values for all tests. As shown in Table 1, LBP and TLX feature sets are not informative to bladder cancer patient stratification as they do not yield significant separation in survival between different prognosis groups. On the other hand, the TIL-related features provide better separations, especially TIL-S and TIL-F-S feature sets. Even though TIL-F or TIL-S alone does not work well when *k*=3, combination of TIL fraction and spatial organization features together stratifies patients into prognosis groups with statistical significance (i.e., *p*=0.039). Fig. 6 shows Kaplan-Meier (KM) plots of TCGA bladder cancer patients stratified into 2 or 3 prognosis groups by using TIL-related image features. For KM plots of patient stratification by other feature sets, please refer to the supplementary file.

**Table 1:** Log-rank test *p*-values on different feature sets. *D*: feature dimensions.

| Feature set | LBP [9]<br>($D = 72$) | TLX [10]<br>($D = 2048$) | TLX [10]<br>($D = 100$) | **TIL-F**<br>($D = 3$) | **TIL-S**<br>($D = 10$) | **TIL-F-S**<br>($D = 13$) |
| --- | --- | --- | --- | --- | --- | --- |
| $k = 2$ | 0.991 | 0.861 | 0.671 | 0.649 | 0.019 | 0.034 |
| $k = 3$ | 0.118 | 0.466 | 0.555 | 0.707 | 0.093 | 0.039 |

**Fig. 6:**
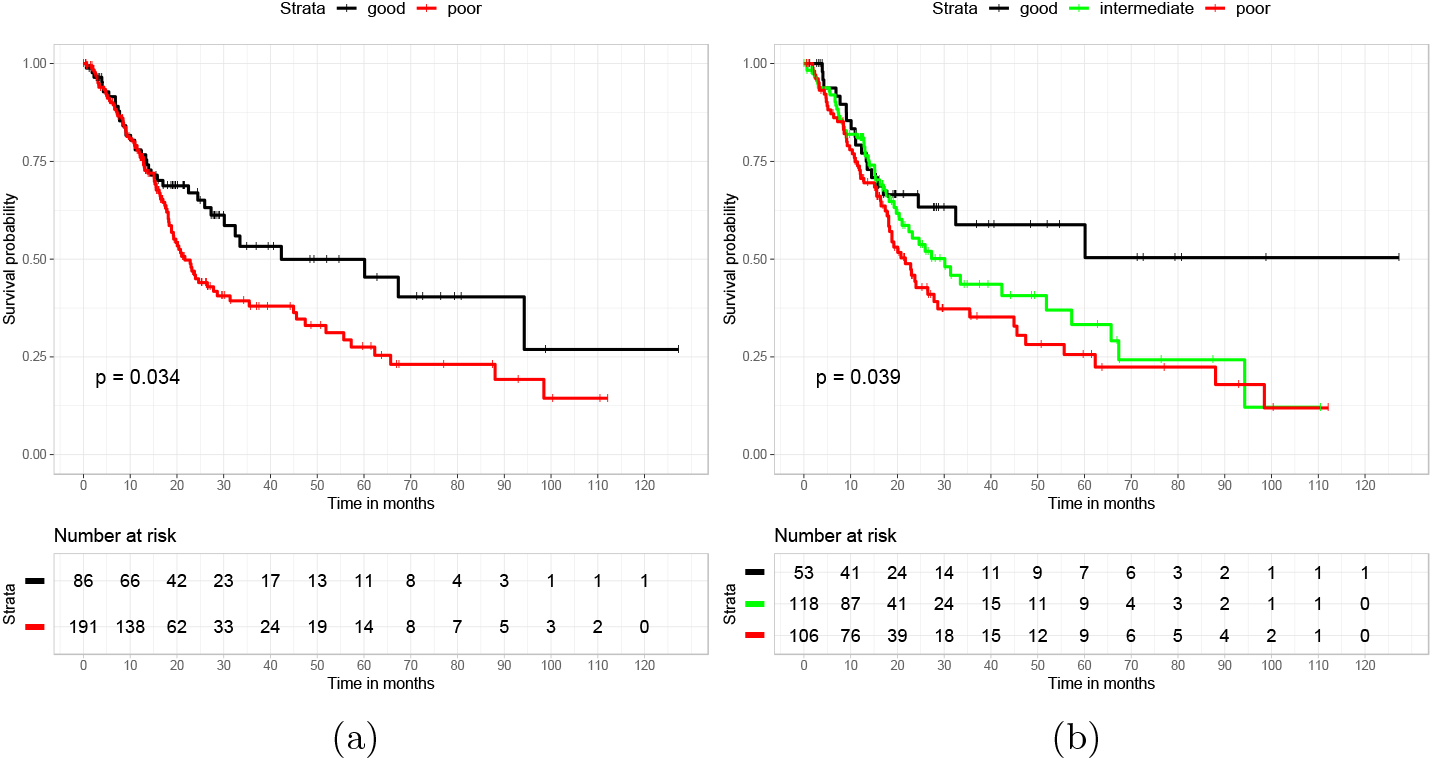
Kaplan-Meier survival curves of bladder cancer patients stratified by fraction and spatial organization of TIL-related image features. (a) *k*=2. (b) k=3.

## 4 Conclusion

We present a study that explores TIL-related histological image features and correlates them with bladder cancer patient survival. Deep CNN models are utilized to detect tumor and TIL regions in the WSI. A novel set of quantitative image features reflecting TIL fraction and spatial organization within tumor region, tumor border region and tumor center region is extracted. Consensus clustering using the features based on TIL fraction and spatial organization is then used to stratify patients into different prognosis groups. Experiments show that quantitative TIL-related image features are informative to stratify TCGA bladder cancer patient with distinct survival outcome. In future, we will attempt to validate our method on other cancer types for patient survival prediction as well as treatment outcomes including immunotherapy.

## References

1. Poch, Michael, et al. “Expansion of tumor infiltrating lymphocytes (TIL) from bladder cancer.” Oncoimmunology 7.9 (2018): e1476816.

2. Saltz, Joel, et al. “Spatial organization and molecular correlation of tumor-infiltrating lymphocytes using deep learning on pathology images.” Cell reports 23.1 (2018): 181–193.

3. Corredor, Germn, et al. “Spatial architecture and arrangement of tumor-infiltrating lymphocytes for predicting likelihood of recurrence in early-stage non-small cell lung cancer.” Clinical cancer research 25.5 (2019): 1526–1534.

4. Simonyan, Karen and Zisserman, Andrew. “Very deep convolutional networks for large-scale image recognition.” arXiv preprint arXiv:1409.1556 (2014).

5. Xu, Hongming, et al. “An efficient technique for nuclei segmentation based on ellipse descriptor analysis and improved seed detection algorithm.” IEEE journal of biomedical and health informatics 18.5 (2014): 1729–1741.

6. Noh, Hyeonwoo, et al. “Learning deconvolution network for semantic segmentation.” Proceedings of the IEEE international conference on computer vision, (2015): 1520–1528.

7. Monti, Stefano, et al. “Consensus clustering: a resampling-based method for class discovery and visualization of gene expression microarray data.” Machine learning 52.1-2 (2003): 91–118.

8. Kim, Jingu and Park, Haesun. “Toward faster nonnegative matrix factorization: A new algorithm and comparisons.” IEEE international conference on data mining, (2008): 353–362.

9. Ojala, Timo, et al. “Multiresolution gray-scale and rotation invariant texture classification with local binary patterns.” IEEE transactions on pattern analysis and machine intelligence 7 (2002): 971–987.

10. Chollet, Franois. “Xception: Deep learning with depthwise separable convolutions.” Proceedings of the IEEE conference on computer vision and pattern recognition, (2017): 1251–1258.

